# Dry Powders for Inhalation Containing Monoclonal Antibodies Made by Thin-Film Freeze-Drying

**DOI:** 10.1101/2021.10.29.466456

**Authors:** Stephanie Hufnagel, Haiyue Xu, Sawittree Sahakijpijarn, Chaeho Moon, Laura Q.M. Chow, Robert O. Williams, Zhengrong Cui

## Abstract

Thin-film freeze-drying (TFFD) is a rapid freezing and then drying technique used to prepare inhalable dry powders from the liquid form for drug delivery to the lungs. We report the preparation of aerosolizable dry powders of monoclonal antibodies (mAbs) by TFFD. We first formulated IgG with lactose/leucine (60:40 w/w) or trehalose/leucine (75:25). IgG 1% (w/w) formulated with lactose/leucine (60:40 w/w) in phosphate buffered saline (PBS) (IgG-1-LL-PBS) and processed by TFFD was found to produce the powder with the most desirable aerosol properties. We then replaced IgG with a specific antibody, anti-programmed cell death protein (anti-PD-1 mAb), to prepare a dry powder (anti-PD1-1-LL-PBS), which performed similarly to the IgG-1-LL-PBS powder. The aerosol properties of anti-PD1-1-LL-PBS were significantly better when TFFD was used to prepare the powder as compared to conventional shelf freeze-drying (shelf FD). The dry powder had a porous structure with nanoaggregates. The dry powder had a Tg value between 39-50 °C. When stored at room temperature, the anti-PD-1 mAb in the TFFD powder was more stable than that of the same formulation stored as a liquid. The addition of polyvinylpyrrolidone (PVP) K40 in the formulation was able to raise the Tg to 152 °C, which is expected to further increase the storage stability of the mAbs. The PD-1 binding activities of the anti-PD-1 mAbs before and after TFFD were not different. While protein loss, likely due to protein binding to glass or plastic vials and the TFF apparatus, was identified, we were able to minimize the loss by increasing mAb content in the powders. Lastly, we show that another mAb, anti-TNF-α, can also be converted to a dry powder with a similar composition by TFFD. We conclude that TFFD can be applied to produce stable aerosolizable dry powders of mAbs for pulmonary delivery.

## Introduction

Protein biologics are stored as liquids or frozen liquids partly due to the relative ease of their manufacture^1^. Liquid storage is not optimal, however, due to poor stability, including aggregation, denaturation, and hydrolysis^2^. This is particularly problematic for large proteins, such as monoclonal antibodies (mAbs), which are expensive to produce and their structure and stability are complex. Antibodies are often provided as intravenous solutions or high concentration liquids in prefilled syringes that must be transported and stored at cooled temperatures (i.e., 2-8°C). Exposure to high or freezing temperatures without the proper stabilizer(s) can exascerbate their instability and degradation. Even if stored at the recommended temperatures, they can still experience accidental freezing/thawing (e.g., in an unevenly cooled refrigerator), leading to protein denaturation and aggregation^3^.

Formulating mAbs and other proteins as dry powders can help address this issue of instability. Powders can be reconstituted for multiple routes of administration with potential advantageous pharmacologic clinical benefits. Anti-PD-1/PDL-1 antibodies are a novel class of immunotherapy approved for their effectiveness across multiple solid tumors administered as intravenous formations. Interestingly, the anti-PD-1 antibody, pembrolizumab (Keytruda®) was originally approved as a powder for reconstitution and intravenous (IV) administration^4^. In addition to reconstitution for IV administration, dry powders that can be aerosolized can be directly delivered to the lungs using a dry powder inhaler (DPI). Many antibodies with respiratory indications are approved by the United States Food and Drug Administration (FDA), with lung cancer being the most common therapeutic indication^5, 6^. Omalizumab (anti-IgE) has been studied in humans for the treatment of asthma via pulmonary delivery but it is currently only approved for subcutaneous (subQ) injection. Typically delivered via injection, E25 (anti-IgE mAb) was aerosolized by nebulization using a PARI IS-2 nebulizer powered by a PARI Master compressor for the treatment of allergen-induced bronchospasm in asthmatic patients^7^. In contrast to IV E25, aerosolized E25 was well-tolerated but was ineffective in controlling symptoms. When delivered via nebulization, E25 was detected in the bronchoalveolar lavage (BAL) and serum of patients. However, the E25 serum concentration was less than 1% of that achieved from IV administration^7, 8^. Therefore, the lack of efficacy was likely due to E25’s low systemic neutralization of IgE, because even if IgE in the lungs is neutralized, the pool of IgE from the blood may redistribute to the lungs^7^. While systemic absorption was necessary for E25’s efficacy in the treatment of asthma, the low systemic absorption of the antibody could be exploited for local therapy in the lungs.

Cetuximab and infliximab have undergone preclinical evaluation for pulmonary delivery. Cetuximab was also formulated as a liquid, while infliximab was formulated as a powder via spray drying (SD)^9^. SD as well as cryogenic techniques including conventional shelf FD, spray-freeze drying (SFD), and spray-freeze into liquid (SFL) and drying are techniques employed to formulate powders^10^. TFFD is a cryogenic technology that can be applied to engineer brittle matrix dry powders of small molecules with good aerosol performance for pulmonary delivery^11–14^. Previously, TFFD was successfully applied to prepare dry powders of proteins such as lysozyme and lactose dehydrogenase (LDH) while preserving their enzymatic activity^10, 15^, but the aerosol properties of the protein dry powders were not reported.

A few classes of mAbs have been approved by the FDA for the treatment of lung cancer: anti-epidermal growth factor receptor (anti-EGFR) mAbs, anti-vascular endothelial growth factor receptor 2 (anti-VEGFR2) mAbs, anti-vascular endothelial growth factor A (anti-VEGF-A) mAbs, and anti-PD-1 mAbs. Guilleminault *et al*. explored the delivery of cetuximab, an anti-EGFR mAb, via aerosolization using a Microsprayer IA-1b aerosolizer introduced orotracheally into the lungs of mice with orthotopic lung tumors^16^. Aerosolization of cetuximab in solution led to a 4-fold higher lung tumor distribution of cetuximab as compared to IV injection at 2 h after dosing^16^. Furthermore, aerosolized cetuximab was able to cause a decrease in mean tumor volume by 37% as compared with saline (p<0.05)^16^, showing that the mAbs retained their function after pulmonary delivery. Similar studies and results were produced by Maillet *et al*^17^. Hervé *et al*. administered G6-31, an anti-VEGF mAb, via aerosolization using a Microsprayer aerosolizer to mice. The plasma C_max_ of G6-31 from the aerosol delivery was about 100-fold less than that of IV delivery, with an estimated bioavailability of 5.1%. Therefore, G6-31 could be used for the treatment of local tumors in the lungs, but likely not systemic metastases or other malignancies. Anti-PD-1 mAbs are unique in that they allow the immune system to kill tumor cells. Unfortunately, they are associated with immune related adverse events (irAEs) that limit their use, and local delivery may improve their tolerability while maintaining their efficacy^18^.

In addition to lung cancer, and potentially solid malignancies with prominent pulmonary metastases, there are other diseases that can benefit from administration of mAbs via the pulmonary route^19^. Anti-tumor necrosis factor alpha (anti-TNF-α) agents have been tested in the clinic for treating idiopathic pulmonary fibrosis^20^ and pulmonary sarcoidosis^21–24^, but outcomes have not been promising due to inconsistent efficacy and relapse^25^. No anti-TNF-α mAbs have been approved to treat pulmonary conditions to date. An anti-TNF-α Fab fragment was delivered intrapleurally, reducing talc-induced pleurodosis in rabbits^26, 27^.

In this work we formulated anti-PD-1 mAbs into dry powders for pulmonary delivery using TFFD and characterized the physicochemical and aerosol properties of the resulting powders. In addition, we showed that anti-TNF-α mAbs can also be formulated using a similar experimental approach.

## Materials and Methods

### Materials

*InVivo*Plus rat IgG2a clone 2A3, *InVivo*MAb anti-mouse PD-1 clone RMP1-14 monoclonal antibodies (mAbs) were from BioXCell (Lebanon, NH). SuperBlock™ blocking buffer in PBS (10 mM), Immunlon 4 HBX 96-well plates and 2,4,6-trinitrobenzene sulfonic acid (TNBSA) were from Thermo Fischer (Waltham, MA). Laemmli 2X buffer, Mini-PROTEAN® TGX™ Precast Gels 4-20%, and Bio-Safe Coomassie G-250 were from Bio-Rad (Hercules, CA). Goat anti-Rat IgG-HRP was from Abcam (Cambridge, MA) and recombinant mouse PD-1 His-tag protein was from R&D Systems (Minneapolis, MN). Glycine, Trizma base, L-leucine, α-lactose monohydrate, D-(+)-trehalose dihydrate, sodium phosphate monobasic dihydrate, sodium phosphate dibasic, PBS + Tween 20 0.05%, phosphate buffered saline (PBS), 3,3’,5,5’-tetramethylbenzidine (TMB), sodium bicarbonate and 2-mercaptoethanol were from Sigma-Aldrich (St. Louis, MO). DriSolv methanol anhydrous was from Mettler Toledo (Columbus, OH) and TB syringes with 21G x 1 needles were from BD (Franklin Lakes, NJ). Aluminum pouches were from IMPAK (Los Angeles, CA) and silica desiccant was from W.A. Hammond Drierite (Xenia, OH). Glass serum vials and plastic low-binding polypropylene cryo vials were from DWK Life Sciences (Millville, NJ). Quali-V®-I hydroxypropyl methylcellulose (HPMC) size 3 capsules were from Qualicaps® (Whitsett, NC). Protein ladder (11-250 kDa) was from New England BioLabs (Ipswich, MA).

### Shelf freeze-drying and thin-film freeze-drying

Samples were prepared by dissolving all components in water or PBS (10 mM, pH 7.4), combined in Eppendorf tubes at the appropriate ratios, and cooled on ice. For TFF, 21G needles on 1 mL syringes were used to apply the samples dropwise onto the rotating drum (150 RPM) of a thin-film freezing device at −100 °C. The frozen films were transferred to 5 mL glass or plastic vials, which were half stoppered with rubber stoppers and stored at −80 °C until lyophilization. A VirTis Advantage bench top tray lyophilizer was used (VirTis, Gardiner, NY). For shelf freeze drying (shelf FD), the shelf and the sample were cooled slowly from RT to −40 °C. For lyophilization, primary drying occurred for 1200 min held at −40 °C, then ramped to 25 °C for 1200 min. Secondary drying occurred at 25 °C for 1200 min. The pressure was held constant at no more than 100 mTorr.

### Evaluation of aerosol performance

Dry powder (2-3 mg) was loaded into an HPMC size 3 capsule (VCaps® Plus, Lonza, Morristown, NJ). The capsule was placed in a Plastiape high resistance RS00 DPI that was then attached to a Next Generation Impactor (NGI, Copley Scientific, Nottingham, UK) device containing pans coated with Tween 20 (1.5% in methanol, w/v), and the methanol was allowed to evaporate prior to initiation of the inhalation. The flow rate was 60 L/min for 4 s per actuation, providing a 4 kPa pressure drop across the device. Each stage was collected with 2 mL water with the exception of the throat, which was collected in 4 mL water. For formulations containing leucine, a TNBSA kit was used to determine the leucine content in each stage. Briefly, samples were diluted with 0.1 M sodium bicarbonate, pH 8.5 and TNBSA 0.01% diluted in methanol and incubated at 37 °C for 50 min. The absorbance was read at 335 nm using a Synergy HT plate reader (BioTek, Winooski, VT). For formulations lacking leucine, the sugar content (i.e. sucrose) was measured using an XBridge Amide 3.5 μm, 4.6 × 150 mm column (Waters, Milford, MA) on a 1220 Infinity II HPLC (Agilent, Santa Clara, CA). The mobile phase was water/acetonitrile 20:80 to 60:40 at a flow rate of 1.0 mL/min for 6 min. The injection volume was 15 μL and the column temperature was 30 °C. A 1290 Infinity II ELSD (Agilent, Santa Clara, CA) was used to detect the sugar with evaporation and nebulization temperatures of 60 °C and a gas flow rate of 1.6 L/min. The Copley Inhaler Testing Data Analysis Software (v3.10) (CITDAS, Copley Scientific, Nottingham, UK) was used to calculate the aerosol properties, including mass median aerodynamic diameter (MMAD), geometric relative standard deviation (GSD), fine particle fraction (FPF) of recovered dose and of delivered dose. The FPF of recovered dose was calculated as the total amount of leucine collected with an aerodynamic diameter below 5 μm as a percentage of the total amount of excipient collected, while the FPF of delivered dose was calculated as the total amount of leucine collected with an aerodynamic diameter below 5 μm as a percentage of the total amount of excipient deposited on the adapter, the induction port, stages 1–7 and Micro-Orifice Collector (MOC).

### Size exclusion chromatography (SEC) and sodium dodecyl sulfate–polyacrylamide gel electrophoresis (SDS-PAGE)

Samples were reconstituted with an equal volume of water as the beginning volume prior to TFFD, bringing the PBS concentration to 1X (10 mM, pH 7.4). A small aliquot of sample was then removed and saved for SDS-PAGE while the remainder of the sample was filtered using a 0.45 μm polyethersulfone (PES) filter. For SEC, an Agilent 300 Å, 2.7 μm, 4.6 × 150 mm column was used on the 1260 Infinity system (Agilent, Santa Clara, CA). The mobile phase was 150 mM sodium phosphate buffer at pH 7, the flow rate was 0.3 mL/min, the time was 10 min, and the wavelength was 220 nm. Unfiltered samples were prepared in laemmli buffer at 1X and beta-mercaptoethanol 10% (v/v) and heated for 5 min at 95 °C. Samples were then loaded into the Mini-PROTEAN® TGX™ Precast Gels 4-20% using the BioRad Tetra cell. The running buffer consisted of 25 mM Trizma, 190 mM glycine, and 0.1% SDS in water. The gel was run for 1 h at 100 V. Afterwards, the gel was washed in distilled water and stained with Bio-Safe Coomassie G-250 Stain. Quantification was conducted using ImageJ.

### Modulated differential scanning calorimetry (mDSC)

A Model Q20 (TA Instruments, New Castle, DE) differential scanning calorimeter equipped with a refrigerated cooling system (RCS40, TA Instruments, New Castle, DE) was used. Powder (3-5 mg) was accurately weighed and loaded into Tzero aluminum hermetic crucibles. A puncture was made in the lid right before the DSC measurement. Samples were first cooled down to −40 °C at a ramp rate of 10 °C/min and then ramped up from −40 to 300 °C at a rate of 5 °C/min. The rate of dry nitrogen gas flow was 50 mL/min. The scans were performed with a modulation period of 60 s and a modulatd amplitude of 1 oC. A TA Instruments Trios v.5.1.1.46572 software was used to analyze the data.

### X-ray powder diffraction (XRPD)

A Rigaku Miniflex 600 II (Rigaku, Woodlands, TX) equipped with primary monochromated radiation (Cu K radiation source, λ = 1.54056 Å) was used. Samples were loaded on the sample holder and analyzed in continuous mode under the operating conditions of accelerating voltage of 40 kV at 15 mA, step size of 0.02° over a 2θ range of 5–40°, scan speed of 1°/min, and dwell time of 2 s.

### Scanning electron microscopy (SEM)

The particle morphology was examined using SEM (Zeiss Supra 40C SEM, Carl Zeiss, Heidenheim an der Brenz, Germany). A small amount of bulk powder (i.e. a flake of TFFD powder) was deposited on the specimen stub using double-stick carbon tape. A sputter was used to coat all samples with 15 mm of 60/40 Pd/Pt before capturing the images.

### Moisture content measurement

A Mettler Toledo V20 volumetric Karl Fischer (KF) titrator (Columbus, OH) was used to measure moisture content in each sample with 5-10 mg of powder diluted into CombiMethanol from Aquastar (Darmstadt, Germany), with 3 independent samples per group.

### Stability study

After lyophilization but prior to removal from the lyophilizer, the samples were backfilled with dry nitrogen gas and the stopper re-cap function was used to cap the vials. Samples were sealed and packaged in aluminum pouches with silica desiccant and the pouches were flushed with nitrogen gas before sealing. They were stored inside a desiccator at the specified temperatures (i.e. 4 °C, room temperature, or 40 ºC). Six or ten weeks later, samples were removed from storage conditions immediately prior to analyzing by SEC, SDS-PAGE, and KF.

### Antibody binding capacity

An enzyme-linked immunosorbent assay (ELISA) was used to measure the binding capacity of the anti-PD-1 mAb. Recombinant mouse PD-1 protein was coated at a concentration of 1 μg/mL on Immunlon 4 HBX plates and incubated overnight at 4 °C. The next day, plates were washed 4 times with PBS + 0.05% Tween 20 (i.e. wash buffer) and blocked with SuperBlock™ buffer according to the manufacturer’s instructions. The plates were washed again and then samples diluted in PBS to a concentration of approximately 100 ng/mL were added and incubated on a plate shaker at room temperature for 2 h. The plate was washed and then IgG-HRP 1:5000 was added, shaking at room temperature for 1 h. The plate was washed and then TMB was added. The reaction was stopped with 0.16 M of sulfuric acid and the plate was read at 450 nm using a Synergy HT plate reader.

### Statistical analysis

Statistical analysis was completed with one-way ANOVA tests with Tukey’s multiple comparisons tests or student’s t tests. A *P* value of ≤0.05 was considered significant. Statistical analyses were performed with GraphPad Prism (San Diego, CA).

## Results and Discussion

### Aerosol performance of thin-film freeze-dried mAb powders

Initially, a non-specific mouse IgG was used as a model antibody to prepare thin-film freeze-dried mAb powders. We previously have identified trehalose/leucine (75:25 w/w) and lactose/leucine (60:40 w/w) as the optimal ratios for formulating thin-film freeze-dried powders with desirable aerosol performance (unpublished data). Therefore, we prepared dry powders of the IgG at a drug loading of 0.5 or 1% (w/w vs. weight of trehalose or lactose plus leucine) and evaluated their aerosol performance. While all formulations exhibited good aerosol performance, increasing the mAb content for the trehalose/leucine-containing formulations from 0.5% to 1% significantly reduced the FPF recovered (Table 1). When IgG dry powders with 1% mAb loading were prepared with lactose/leucine, however, the FPF and all other aerosol properties were retained at optimal levels (Table 1). We also investigated the use of water versus PBS as the solvent for TFFD. Compared to formulations prepared in water, formulations prepared in PBS showed improved aerosol properties and generally had smaller variability between samples (Table 1). Reproducibility is critical in respiratory delivery, since delivery with DPI can have high patient-to-patient variability. Because of the excellent aerosol performance of the dry powders prepared with lactose and leucine and the fact that lactose is currently the only sugar approved by the FDA as a carrier for pulmonary delivery, the dry powder prepared with 1% IgG and lactose/leucine in PBS (i.e. IgG-1-LL-PBS) was chosen for further studies, with our understanding that lactose is a reducing sugar and may negatively affect the chemical stability of the mAbs during long-term storage. We are also aware that high concentration of salts, especially those in PBS, are known to negatively affect biologics during freezing and drying^28–30^ and understand that for different mAbs and mAb contents, one may need to avoid PBS in the mAb solution before subjecting it to TFFD to better preserve the integrity of the mAbs.

**Table 1.** Composition and aerosol performance properties of IgG powders prepared by TFFD. TL = trehalose and leucine; LL = lactose and leucine. Data are mean ± S.D. (n = 3).

| Formulation | Lactose mAb loading (% w/w) | Ratio | | | Solid content (% w/v) | MMAD (μM) | GSD | FPF (% recovered) | FPF (% delivered) |
| --- | --- | --- | --- | --- | --- | --- | --- | --- | --- |
|  |  | Lactose | Trehalose | Leucine |  |  |  |  |  |
| IgG-0.5-TL | 0.5 | - | 75 | 25 | 1 | 2.438 ± 0.941 | 3.507 ± 0.588 | 67.335 ± 11.530 | 71.225 ± 12.727 |
| IgG-0.5-TL-PBS | 0.5 | - | 75 | 25 | 1 | 1.671 ± 0.129 | 2.732 ± 0.169 | 78.588 ± 4.904 | 85.453 ± 3.342 |
| IgG-1-TL | 1 | - | 75 | 25 | 1 | 3.161 ± 0.222 | 2.118 ± 0.044 | 46.982 ± 12.077 | 59.830 ± 9.541 |
| IgG-1-TL-PBS | 1 | - | 75 | 25 | 1 | 2.145 ± 0.207 | 2.048 ± 0.278 | 48.252 ± 5.225 | 82.128 ± 11.780 |
| IgG-1-LL | 1 | 60 | - | 40 | 1 | 2.578 ± 0.389 | 1.871 ± 0.112 | 66.729 ± 16.502 | 82.247 ± 12.555 |
| IgG-1-LL-PBS | 1 | 60 | - | 40 | 1 | 1.693 ± 0.064 | 1.863 ± 0.073 | 92.636 ± 1.311 | 97.504 ± 1.257 |

We then prepared anti-PD-1 mAb dry powders (anti-PD1-1-LL-PBS) using the composition similar to the IgG-1-LL-PBS by TFFD or shelf FD and evaluated their aerosol properties. As shown in Figs. 1A-B, the TFFD powder demonstrated significantly better aerosol performance as compared to the dry powder with identical composition but prepared by shelf FD, with a smaller MMAD value and larger FPF values (Fig. 1B). Shelf FD is known to produce powders that have relatively low surface area, which contribute to poor aerosol performance^10^. This is in large part due to the low freezing rate, which allows protein particles to grow over a longer time to reach the frozen state. Conversely, TFFD has a higher freezing rate, which, in addition to the shorter time to reach the frozen state allowing for less particle growth, also leads to more nucleated ice domains with thinner liquid channels. Thinner channels correspond to fewer particle collisions and less particle growth, yielding smaller particles adequate for pulmonary delivery^15^. SD is another popular technique for producing powders, but exposes the formulation to heat, which can be particularly damaging for biologics, such as mAbs. The powders resulting from SD processing are dense and, thus, often cannot be easily aerosolized^31^. On the other hand, TFFD powders tend to be porous and have low density, corresponding to readily dispersible powders with high FPFs^13, 14, 32, 33^.

**Figure 1.**
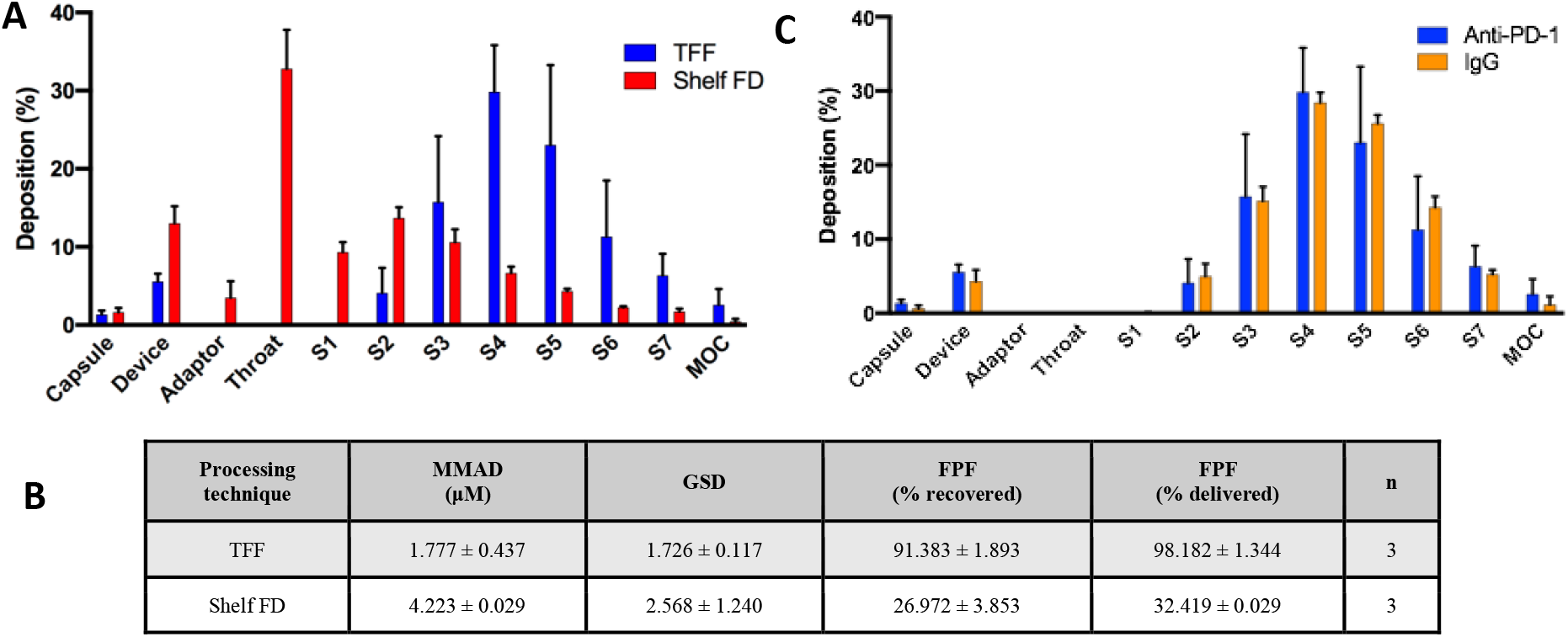
(A-B) Aerosol performance properties of anti-PD1-1-LL-PBS TFFD powder (TFF) compared to anti-PD1-1-LL-PBS shelf FD powder (Shelf FD). (C) Aerosol performance properties of anti-PD1-1-LL-PBS TFFD powder compared to IgG-1-LL-PBS TFFD powder. In B, data are mean ± S.D.

Additionally, applying TFFD to anti-PD-1 mAbs did not significantly affect the aerosol properties of the resultant dry powders (i.e. anti-PD1-1-LL-PBS vs. IgG-1-LL-PBS (Fig. 1B vs Table 1, and 1C)), indicating that the TFFD technology can be applied to prepare aerosolizable dry powders of different mAbs.

### Characterization of the physical properties of the thin-film freeze-dried anti-PD-1 mAb powder

SEM shows that the thin-film freeze-dried anti-PD-1 mAb powder (i.e. anti-PD1-1-LL-PBS) was highly porous and consisted of nanoaggregates (Fig. 2A), which is in alignment with other dry powders produced via TFFD^10^. The moisture content of the anti-PD1-1-LL-PBS powder was 1.5 ± 0.2%. XRPD revealed that lactose was amorphous, while leucine and PBS were crystalline (Fig. 2B). When assessing the physical properties with mDSC, no melting points (MP) for the crystalline PBS could be seen (Fig. 2C). The MPs for sodium chloride and sodium phosphate are >1000 °C and thus were likely out of range^34^. The mDSC of leucine and PBS showed that leucine’s MP was around 260 °C, decreased from its normal melting point of 300 °C. When lactose was combined only with PBS, a T_g_ signal was seen at 128 °C, similar to a typical T_g_ of lactose^35–37^. An MP was also present for lactose at 192 °C, substantially lower than the typical lactose MP of 220°C^38^, which was possibly due to melting point depression by the reduction of the particle size by TFFD, and also showed that lactose in the TFFD powder may be partially crystalline. The lactose T_g_ was still present when leucine was added, but the MP was observed at 140 °C. Upon addition of anti-PD-1 mAb, the T_g_ was reduced to around 51 °C (a value of 39 °C was observed in one test). Additionally, two MPs were present at 134 and 144 °C. According to Gombas et al, and Lopez-Pablos et al., the melting peaks in the range of 130-160 °C that were observed in lactose samples are related to the evaporation of water chemically bonded to the lactose molecule ^39, 40^.

**Figure 2.**
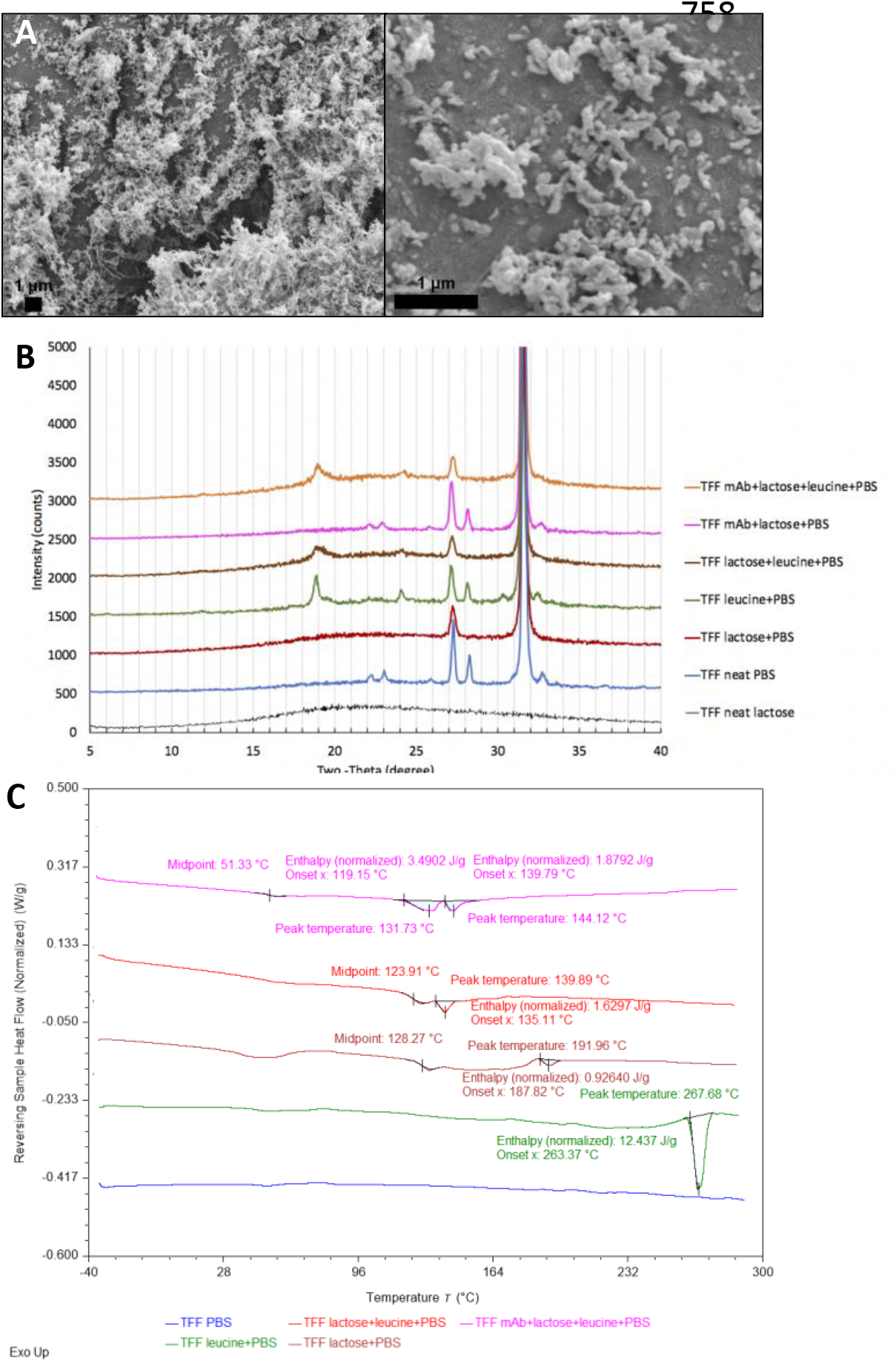
Characterization of anti-PD1-1-LL-PBS TFFD powder by (A) SEM, (B) XRPD, and (C) mDSC.

### Stability of anti-PD-1 mAbs in the thin-film freeze-dried powder

To test the stability of the anti-PD-1 mAbs in the anti-PD1-1-LL-PBS powder, the powder was stored at 4 °C, room temperature, or 40 °C for 10 weeks and the integrity of the mAbs was evaluated using SDS-PAGE and SEC. As a control, the stability of anti-PD-1 mAbs in liquid were also tested. As shown in Figs. 3A-D, the anti-PD-1 mAbs in the thin-film freeze-dried powder were more stable than in the liquid sample at room temperature and 40 °C. However, at 40 °C, the powder showed a slight upward shift in both bands in the SDS-PAGE image (Fig. 3A-B), which was also seen as a left shift in SEC peaks (not shown). SEC data revealed that the percent of monomer was lower in the liquid sample stored at room temperature or 40 °C (Fig. 3C) and indicated that the loss was mostly due to degradation. However, the loss of monomer in dry powder stored at 40 °C was more due to aggregation (Fig. 3C), potentially contributing to the increase in protein size in SDS-PAGE and SEC data (Fig. 3A-B). Both 6- and 10-week stability data showed similar trends in the amount of monomer, with the monomer content decreasing as the storage temperature increased (Fig. 3C). This decrease in monomer was significant in liquid samples at room temperature and 40 °C compared to time 0 (p < 0.05 for both temperatures at 6 and 10 weeks). In contrast, loss of monomer in the dry powder was seen only when stored at 40 °C and was to a smaller degree than liquid samples, not when stored at 4 °C or room temperature (Fig. 3C). Overall, it appears that the anti-PD-1 mAbs in the dry powder was stable after 10 weeks of storage at room temperature, but not at 40 °C.

**Figure 3.**
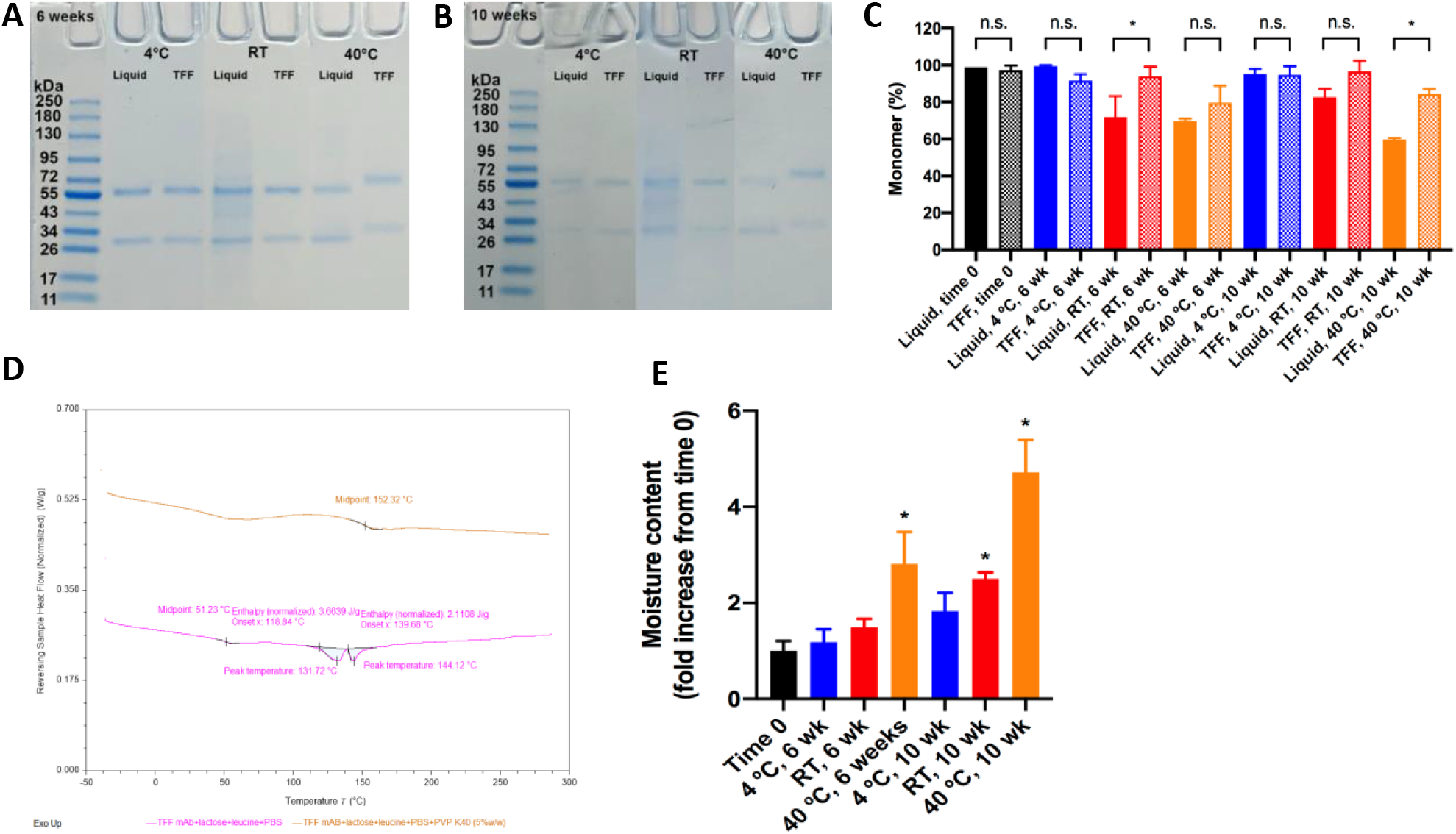
Stability of anti-PD-1 mAbs in liquid or in anti-PD1-LL-PBS dry powder prepared by TFFD. Samples were stored for (A) 6 or (B) 10 weeks and removed from the storage conditions immediately prior to analysis using SDS-PAGE or (C) SEC. (D) mDSC of anti-PD1-1-LL-PBS TFFD powder containing PVP K40. (E) (Relative) moisture content in anti-PD1-1-LL-PBS TFFD powders after 6 or 10 weeks of storage at 4°C, room temperature (RT), or 40°C. For C and E, data are mean ± SD, * indicates significance, and n.s. indicates no significance.

The instability of the anti-PD-1 mAbs in the dry powder stored at 40 °C was not unexpected, however, because the T_g_ of the anti-PD1-1-LL-PBS powder was found to be between 39-51 °C (Fig. 2C). To further improve the thermal stability of the anti-PD-1-LL-PBS dry powder in the future, PVP K40 may be used, as a thin-film freeze-dried anti-PD-1 mAb powder prepared with 5% (w/w) of PVP K40 (i.e. anti-PD1-1-LL-PBS-PVP) showed a T_g_ of 152 °C (Fig. 3D). The ability of PVP K40 to impact the stability and aerosol performance of TFFD mAb formulations will be investigated in the future. To further understand the instability observed when the dry powder was stored at 40 °C, the moisture content in the dry powder samples after storage was also measured. Data showed that anti-PD1-1-LL-PBS powder increased moisture sorption after 6 and 10 weeks of storage at 40 °C (Fig. 3E). Storing the dry powder at room temperature significantly increased the moisture content only after 10 weeks of storage (Fig. 3E). Humidity control and storage conditions are thus important with this hygroscopic powder. Handling of the dry powder in a lower humidity condition and better packaging to minimize moisture uptake during storage are expected to help improve the long-term storage stability of the dry powder without refrigeration or freezing. Nonetheless, data in Fig. 3 suggest the potential of applying the TFFD technology to enable cold chain-free storage of mAbs.

### Binding capacity of anti-PD-1 mAb to PD-1 after subjected to TFFD

The binding capacity of the anti-PD-1 mAbs was measured before and after subjected to TFFD and reconstitution. The anti-PD1-1-LL-PBS powder was resuspended in an equal volume of water as the liquid sample (before TFFD). The binding capacity of the anti-PD-1 mAbs after subjected to TFFD was not statistically different from that before TFFD, though the variability was high after TFFD (Fig. 4A, glass vial). The high variability may be in part due to the interaction of the mAbs with the glass vial, as changing the glass vial to plastic vial led to a significant reduction in the binding capacity of the anti-PD-1 mAbs without significantly affecting the mAb recovery rate (Fig. 4B).

**Figure 4.**
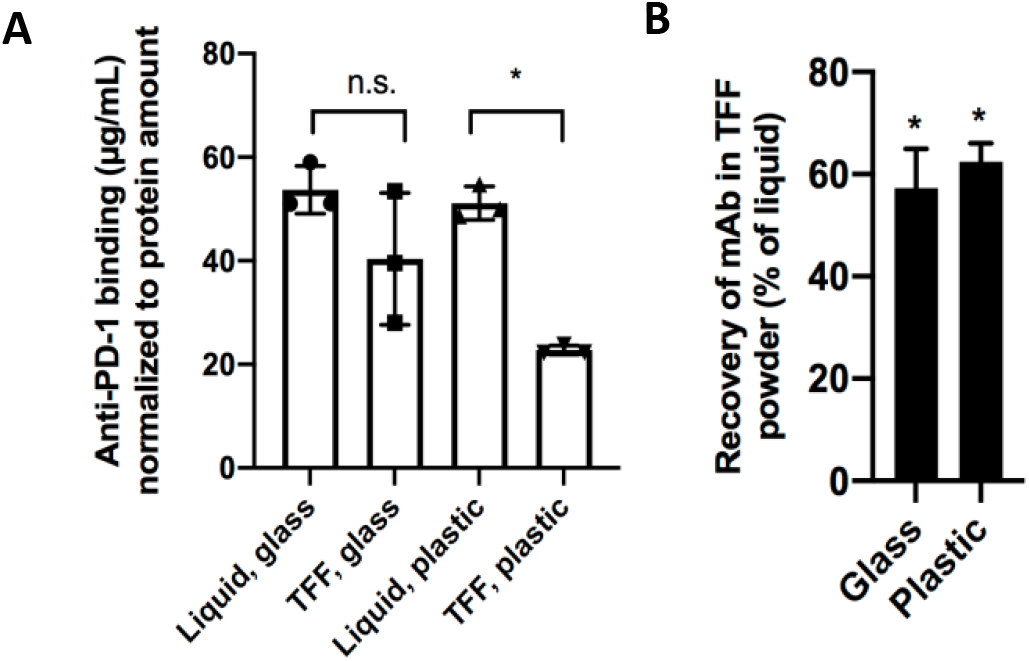
Antigen binding capacity of anti-PD-1 mAbs before and after TFFD. (A) ELISA data normalized to protein content and (B) quantification of protein loss based on SDS-PAGE. Data are mean ± SD, * indicates significance compared to liquid anti-PD-1 mAbs, and n.s. indicates no significance.

### Assessment of reasons underlying protein loss during TFFD

Because the mAb recovery rate of not 100% after TFFD (Fig. 4B), we investigated potential reasons underlying the loss of mAbs during the TFFD. A major consideration in preparing dry powder formulations of proteins is aggregation. Aggregation can lead to immunogenicity that can be detrimental to patient health and must be minimized^41^. We assessed whether TFFD processing affected the degree of aggregation of anti-PD-1 mAb. Both SDS-PAGE and SEC showed no increase in aggregation in mAbs reconstituted from TFFD powder as compared to the same mAb formulation in the liquid state before TFFD (data not show), which prompted us to hypothesize that adsorption of mAbs to various surfaces such as the stainless-steel drum surface of the TFF apparatus, glass vial, plastic disposables. To test this hypothesis, we thin-film freeze-dried the anti-PD-1 mAbs at increased mAbs loading (i.e. mAbs to lactose/leucine at 2.6%, 6.6%, and 13.2%, while maintaining the content and concentration of other ingredients in the mAb solution before subjected to TFFD). As shown in Figs. 5A-B, increasing the mAb concentration in the mAb solution before subjected to TFFD significantly improved the recovery of the mAbs (to close to 100%), indicating that the mAb loss was likely due to surface adsorption during the TFFD process and the adsorption was saturable (Figs. 5A-B), rather than aggregation because it is known that for antibodies, increasing their concentration can often reduce their stability during freeze-drying^42^. The percentage of monomer in the anti-PD-1 mAb TFFD powder prepared at 6.6% and 13.2% of mAbs remained at 96.1 and 96.4%, respectively, although slightly decreased as compared to its liquid counterpart (Fig. 5C). Of note, some aggregation and degradation were visible in the research-grade anti-PD-1 mAb liquid sample. Using a high purity clinical grade mAb in the future should be helpful.

**Figure 5.**
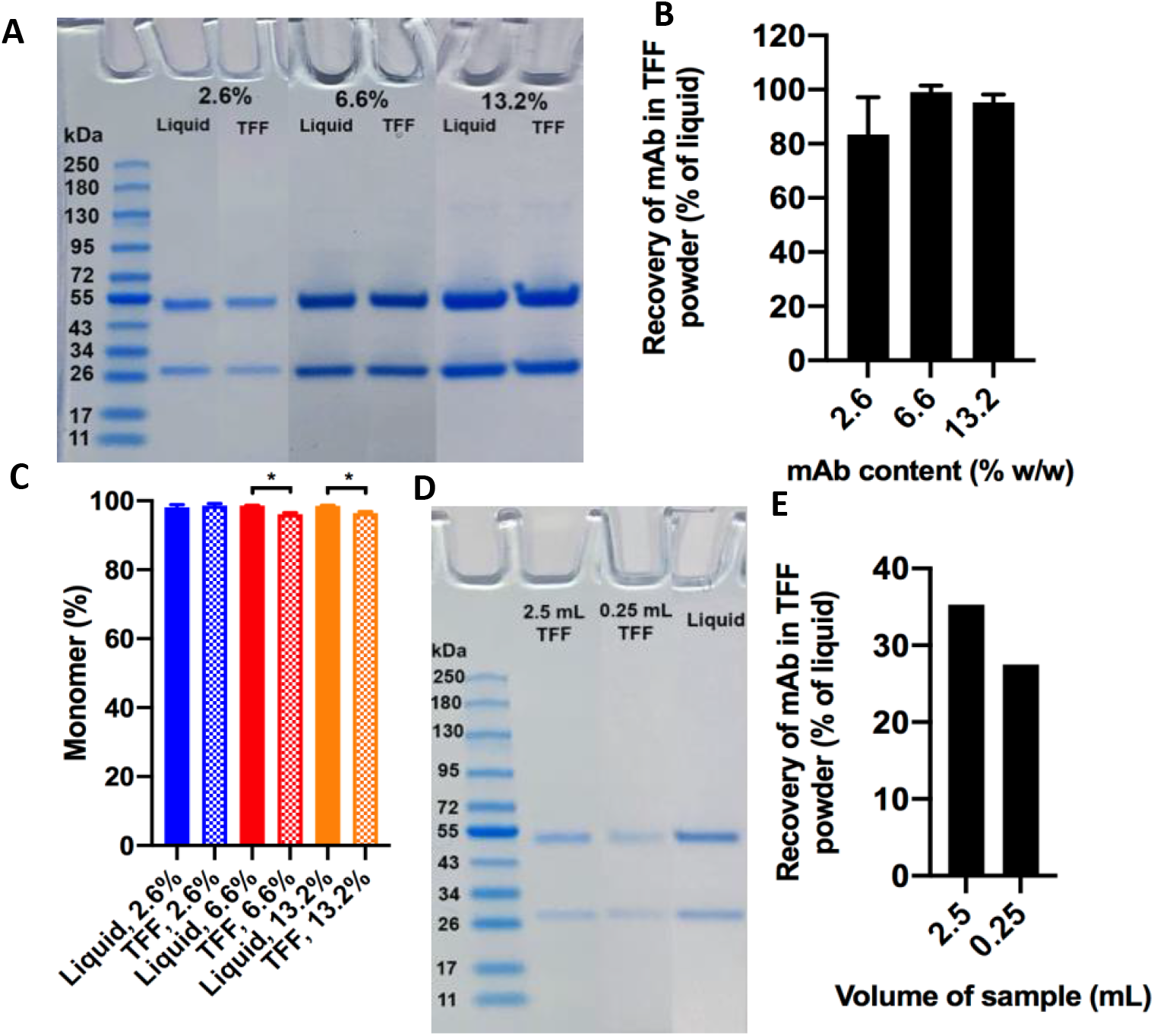
Protein loss saturability. (A) SDS-PAGE of anti-PD-1 mAbs reconstituted from TFFD powders prepared with higher contents of anti-PD-1 mAbs (2.6, 6.6, or 13.2%, w/w) using lactose/leucine (60:40) as the excipients. (B) Quantification of band intensity on SDS-PAGE from A. (C) Monomer content in anti-PD-1 mAbs reconstituted from TFFD powders prepared with higher contents of anti-PD-1 mAbs using lactose/leucine as the excipients. (D) SDS-PAGE of anti-PD-1 mAbs reconstituted from TFFD powders prepared with anti-PD-1 mAbs in a solution containing lactose/leucine as the excipients at different volumes. (F) Quantification of band intensity on SDS-PAGE from D. In B, C, and E, data are mean ± SD, * indicates significance, and n.s. indicates no significance

To further confirm that the mAb loss during TFFD was related to protein adsorption, we thin-film freeze-dried anti-PD-1 mAbs at 1% mAb loading, but in a larger volume (i.e. 0.25 vs. 2.5 mL). As shown in Figs. 5D-E, increasing the volume of the mAb solution for TFFD only slightly reduced protein loss, not as effective as increasing the concentration of mAbs (Fig. 5B vs. Fig. 5E). This makes sense, as increasing the volume alone did not saturate the surface area the mAbs were exposed to during thin-film freezing because the volume and surface area both increased proportionally (i.e. a higher volume of mAb solution corresponds to more thin-films were frozen). This is in contrast to increasing the mAb concentration, in which case the surface area the mAbs were exposed to stayed the same, further supporting that the protein loss observed in the anti-PD1-1-LL-PBS sample was largely due to binding of the mAbs to surfaces during the TFFD process.

### Alternative excipients and mAb recovery

Lactose was largely chosen as the primary excipient when thin-film freeze-drying the anti-PD-1 mAbs in the studies above. Other excipients such as trehalose and sucrose are also being explored for pulmonary delivery although they are not yet in any FDA-approved products for this route. Leucine was included in the powder formulations because data from our previous studies showed that it helps improve the aerosol performance of the TFFD powders^11, 43^. Leucine is also in an inhaled product in clinical trials^44^. The solubility of leucine in water is ~25 g/L at room temperature^45^. Leucine crystalizes during TFFD, and crystalline forms of molecules are known to be less soluble, which may lead to formulation of visible and subvisible particles during TFFD and reconstitution of the mAbs. In fact, when reconstituting the anti-PD1-1-LL-PBS, some visible particles could be seen. Therefore, we also tested i) the effect of using trehalose or sucrose as the primary excipient, with or without leucine to prepare thin-film freeze-dried mAb powders, and ii) their effect on mAb recovery after TFFD. The visible particles were minimized or eliminated when leucine was not included (Fig. 6A) and thus may be due to leucine’s low solubility or dissolution when in the crystalline state. Keytruda’s package insert states to “discard reconstituted vial if extraneous particulate matter other than translucent to white proteinaceous particles is observed,” implying that some visible protein particles are acceptable even for intravenous administration^4^. This formulation has since been discontinued, however. Trehalose/leucine 75:25 (TL) 1% (w/v) and sucrose 5% (w/v) without leucine (S) were also used to formulate anti-PD-1 mAb 1% (w/w) in PBS (anti-PD1-1-TL-PBS and anti-PD1-1-S-PBS) and the protein recovery and percent of monomer were measured. Clearly, excipients impacted the degree of recovery of protein (Figs. 6B-C). Sucrose as an excipient actually improved the protein recovery of the TFFD samples compared to the liquid samples. Additionally, trehalose/leucine had better recovery of protein than lactose/leucine (Figs. 6B-C). Furthermore, sucrose as an excipient did not decrease the percent of monomer in the TFFD powder, while trehalose/leucine did (Fig. 6D). Due to the excellent protein and monomer recovery in the sucrose formulation (Figs. 6A-D), the binding activity and aerosol performance were measured as well. The sucrose formulation maintained similar anti-PD-1 mAb binding before and after processing with TFFD and reconstitution (Fig. 6E). Furthermore, the sucrose formulation had good aerosol performance even without leucine (Fig. 6D). It is hypothesized that leucine could be responsible in part to protein loss, with 0% leucine having the best recovery (Figs. 6B-C) as well as no visible aggregates upon reconstitution (Fig. 6A). Therefore, future formulation efforts will be focused on identifying the proper leucine content in the powder so that the protein loss will be minimized, while the resultant dry powder still has good aerosol performance properties.

**Figure 6.**
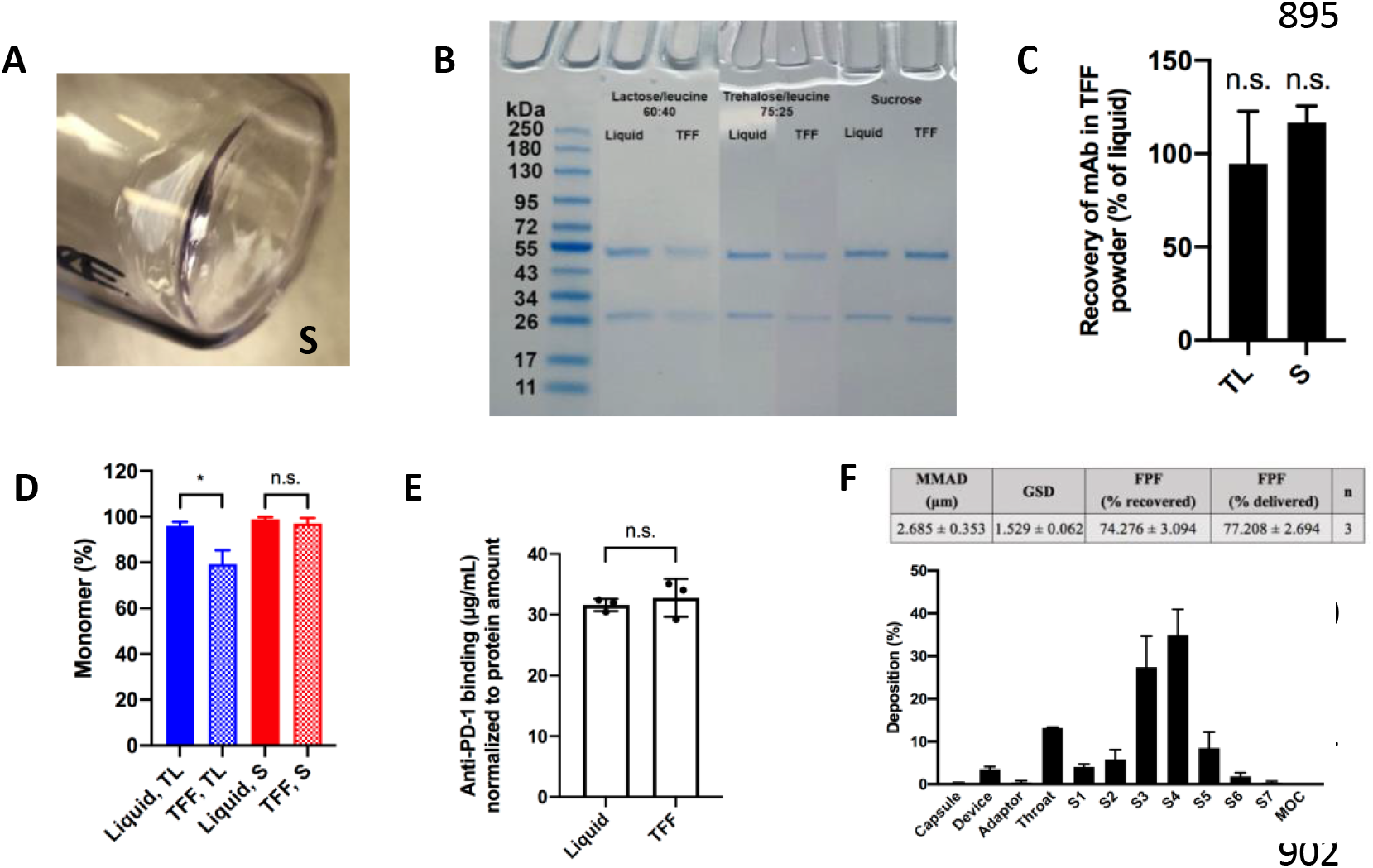
Excipient choice impact on mAb recovery after TFFD. (A) Lack of visible aggregates in anti-PD-1 mAb TFFD powder prepared with sucrose alone as the excipient. (B) SDS-PAGE comparing anti-PD-1 mAbs reconstituted from three powders of different compositions (lactose/leucine, 60:40; trehalose/leucine (TL), 75:25; or sucrose (S) alone). (C) Quantification of protein from SDS-PAGE for TL and S. (D) Percent of monomer in anti-PD-1 mAb TFFD powders prepared with trehalose/leucine (75:25) (TL) or sucrose alone (S) as determined by SEC. (E) ELISA data showing the PD-1 binding activity of the anti-PD-1 mAbs reconstituted from the anti-PD-1 mAb TFFD powder prepared with sucrose alone as the excipient. Data were normalized to protein content. (F) Aerosol properties anti-PD-1 mAb TFFD powder prepared with sucrose alone as the excipient. In C-F, data are mean ± SD (n = 3).

### Thin-film freeze-drying of anti-TNF-a mAb

Finally, anti-TNF-α mAb was also formulated at 1% (w/w) with trehalose/leucine 75:25 in PBS (anti-TNFα-1-TL-PBS). Interesting, there was a full recovery of the protein (Figs. 7A-B). When anti-PD-1 mAb was formulated at 1% with trehalose/leucine 75:25, the monomer content decreased from 96.0 to 79.2% after TFFD (Fig. 6C). However, when the mAb was changed to anti-TNF-α mAb while other formulation parameters were the same, the monomer percentage did not decrease after TFFD (Fig. 7C). Therefore, as expected, the characteristics of each mAb, including the composition of the liquid in which the mAb is dissolved, can also impact the recovery and stability of the protein during TFFD and each mAb requires its own formulation optimization.

**Figure 7.**
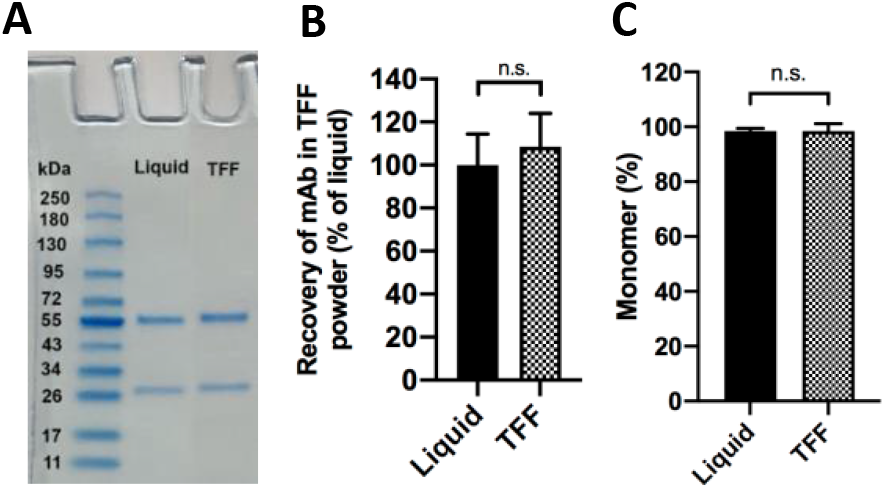
Characterization of anti-TNFα mAb dry powder prepared by TFFD. (A) SDS-PAGE showing anti-TNFα mAbs before and after subjected to TFFD and reconstitution. (B) Quantification of protein from SDS-PAGE gel in A. (C) Percent of anti-TNFa mAb monomer before and after subjected to TFFD and reconstitution.

## Conclusion

TFFD can be utilized to prepare mAbs as stable powders with desirable aerosol performance properties for pulmonary delivery. The excipient type, concentration and properties, as well as the specific type of mAb and its concentration can impact the protein recovery and monomer content in the TFFD powders and should be optimized for each mAb. Additionally, moisture control must be maintained to maximize stability of the mAbs when stored at high temperatures, and polymers such as PVP can be used to modify the T_g_ and thus can increase the physical stability of the mAbs contained in the TFFD powders. The mAb dry powders may have potential beneficial effects if explored clinically in the treatment of cancer.

## Acknowledgements

This work was in part supported by TFF Pharmaceuticals, Inc. (to ROW and ZC), UT Austin Dell Medical School Texas Health Catalyst (to ZC), and the Alfred and Dorothy Mannino Fellowship in Pharmacy (to ZC).

